# Reversions to consensus are positively selected in HIV-1 and bias substitution rate estimates

**DOI:** 10.1101/2022.02.13.480259

**Authors:** Valentin Druelle, Richard A. Neher

## Abstract

HIV-1 is a rapidly evolving virus able to evade host immunity through rapid adaptation during chronic infection. The HIV-1 group M has diversified since its zoonosis into several subtypes at a rate of the order of 10^−3^ changes per site per year. This rate varies between different parts of the genome and its inference is sensitive to the time scale and diversity spanned by the sequence data used. Higher rates are estimated on short time scales and particularly for within-host evolution, while rate estimates spanning decades or the entire HIV-1 pandemic tend to be lower. The underlying causes of this difference are not well understood.

We investigate here the role of rapid reversions toward a preferred evolutionary sequence state on multiple time scales. We show that within-host reversion mutations are under positive selection and contribute substantially to sequence turnover, especially at conserved sites. We then use the rates of reversions and non-reversions estimated from longitudinal within-host data to parametrize a phylogenetic sequence evolution model. Sequence simulation of this model on HIV-1 phylogenies reproduces diversity and apparent evolutionary rates of HIV-1 in *gag* and *pol*, suggesting that a tendency to rapidly revert to a consensus-like state can explain much of the time dependence of evolutionary rate estimates in HIV-1.

## Introduction

RNA viruses have low fidelity polymerases, resulting in a rapidly diversifying virus population, which, in turn, facilitates the adaptation to changing environments. The human immunodeficiency virus 1 (HIV-1) is a prime example of such a rapidly evolving virus. The life-long infections it causes are characterized by a large viral population that accumulates diversity at a high rate to constantly evade host immunity (Coffin, Swanstrom, 2013). This continuous evolution has led to a diverse viral population on the pandemic scale that is categorized into several viral subtypes (Brian Foley, 2018; Li et al., 2015). Different lineages have accumulated diversity at a rate of about 1 substitution in 1000 sites per year since its jump to human hosts at the turn of the 20th century (McCutchan, 2006; Sharp, Hahn, 2011; Korber et al., 2000).

Quantifying the rate of viral evolution, however, is surprisingly difficult and different approaches yield different answers. Most importantly, the time scale across which sequences are compared affects the estimates strongly, sometimes by orders of magnitude: the longer the time scale, the lower the estimate (Aiewsakun, Katzourakis, 2016; Hanada et al., 2004; Worobey et al., 2010; Gilbert, Feschotte, 2010; Ghafari et al., 2021). These discrepancies suggest that we lack a good understanding of how micro-evolutionary within-host (WH) processes on the scales of days, months and years give rise to the diversity observed on longer time scales across hosts. In the case of chronic infections such as HIV-1, these micro-evolutionary processes are driven by selection to evade a changing host immune response at each transmission while maintaining fitness.

HIV-1 is an ideal system to study these effects in detail since the rate discrepancies between the WH, pandemic and broader scales are well documented (Alizon, Fraser, 2013; Worobey et al., 2010), the pandemic is well sampled, and high resolution WH data exists. The evolutionary rate estimated on the pandemic scale is around two to five times lower than the one observed within host (Alizon, Fraser, 2013). Several hypotheses have been put forward to explain this phenomenon. Two of the main hypotheses are the preferential transmission of ancestral HIV-1 variants, i.e. the “store and retrieve” hypothesis (Lythgoe, Fraser, 2012), and the reversion toward an ancestral-like state, i.e. the “adapt and revert” hypothesis (Redd et al., 2012; Zanini et al., 2015; Leslie et al., 2004). The relative importance of these and possibly other processes for the discrepancy of rate estimates is not well understood (Raghwani et al., 2018).

We use WH longitudinal deep sequencing data to investigate the evolutionary processes responsible for the difference of evolution rate observed. We, firstly, show that rate estimates are related to the time scale and reference point used on both the WH and pandemic scale, suggesting a saturation of divergence long before it is expected in most sequence evolution models. Secondly, we investigate the cause of this saturation and find that reversion towards the HIV-1 consensus is more common than expected. Such WH reversion mutations are positively selected. Lastly, we use simulations of evolution to quantify the impact of such reversions on rate estimates on time scales of decades and above. More generally, our results highlight the evolutionary bias of viruses toward a consensus or state of high intrinsic fitness in a changing environment.

## Results

We use a (i) a set of sequences representative of the HIV-1 pandemic spanning multiple decades and a (ii) longitudinal data set following the evolution of the virus within individual hosts to investigate patterns of evolution on multiple time scales. The former between-host (BH) data set contains 1000 HIV-1 group M sequences from the Los Alamos National Laboratory HIV database (Foley et al., 2013). Subsampling was performed to have the same number of sequences for each year to avoid sampling biases (except for early years, where fewer sequences are available) but otherwise randomly picked from the full data set. Our WH analysis is based on the HIVEVO data set (Zanini et al., 2015), a whole genome deep sequencing of HIV-1 populations in 11 patients during a 4 to 16 years follow-up without treatment. Between 6 and 12 samples are available per patient. Sequencing depth and template input of all samples in this data set has been assessed and most samples allow a confident calling of frequencies of minor variation down to a few percent (Zanini et al., 2016). See section M&M 1.1 for details.

We analyze the evolution of the *env, pol* and *gag* genes of HIV-1 below. They code for surface proteins, viral enzymes and capsid proteins, respectively (Freed, 2001). When combined, they cover approximately 80% of the genome. We focus on the *pol* region in the main text and present analogous results for the *env* and *gag* regions in the Supplementary Materials.

### Apparent rates depend strongly on the distance to the reference sequences

The rate at which differences between sequences accumulate decreases with time as more and more sites are hit multiple times by changes (Felsenstein, 2004). If all sites evolve at the same speed, such saturation effects are only important once distances between sequences are large (the size of correction is proportional to the distance squared). However, if different sites evolve at drastically different rates, or reversions are common, such saturation effects set in much earlier and can lead to significant deviations even when sequences are still very similar (Puller et al., 2020; Ghafari et al., 2021).

Figure 1 compares the rate of accumulation of sequence changes of HIV-1 at the pandemic scale (A) and within hosts (B) in the *pol* gene. Figure 1A shows the Hamming distance of HIV-1 sequences from the inferred root of the HIV-1 group M tree (orange) or the consensus of the subtype (green). These Hamming distances are compared to the corresponding increase in root-to-tip (RTT) differences along a reconstructed phylogeny (blue).

**Figure 1:**
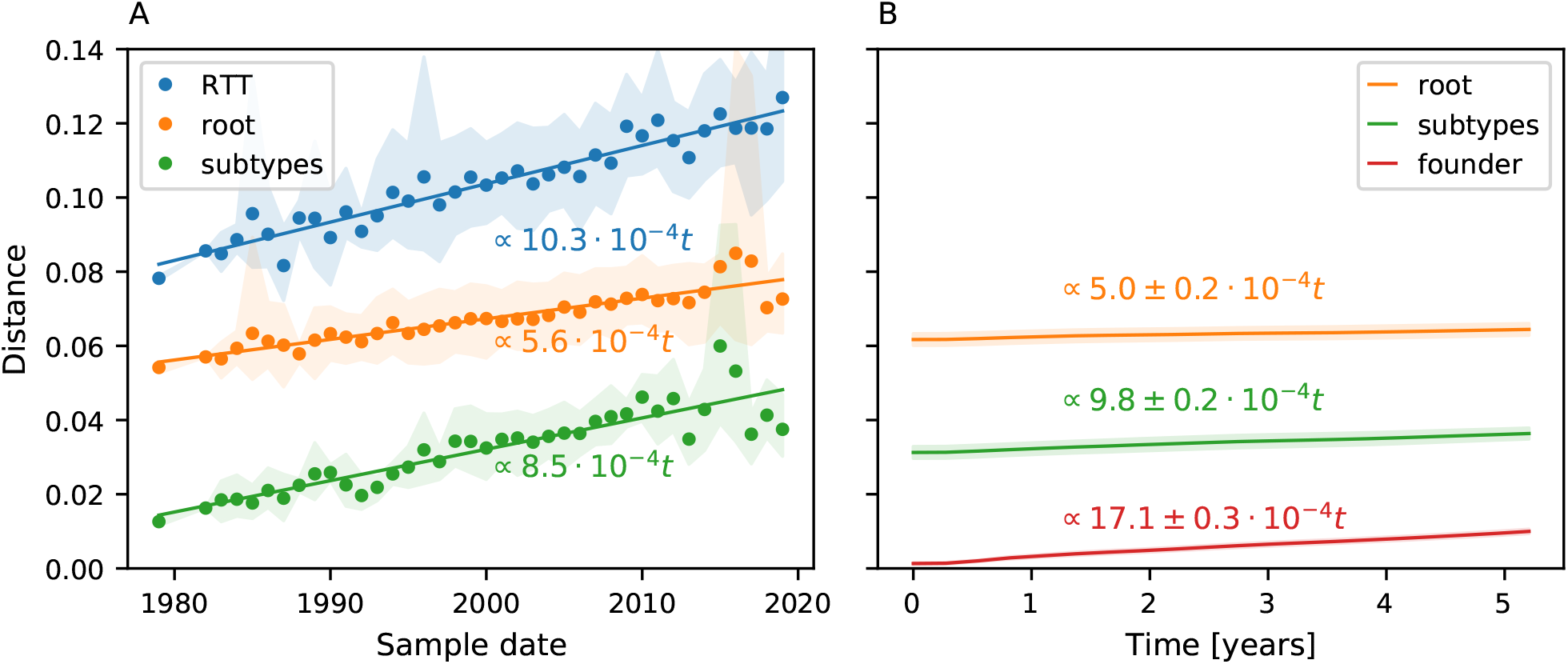
Divergence over time in the *pol* gene. **A**: The graph shows the average Hamming distance from the root of the HIV-1 group M tree, from the respective subtype, or RTT distance as a function of time. Each data point is the average of sequences from one year; the shaded area indicates the 10-90% range. **B**: The WH divergence over time relative to the putative founder genotype and HIV-1 group M root or subtype consensi, averaged over all patients in the HIVEVO data set. Divergence is computed according to equations 1 and 2. The Hamming distance to the HIV-1 group M root or the subtype consensi increases at similar rates within hosts and cross-sectionally. Results for region *env* and *gag* are shown in Supplementary Figures S1 and S2.

The slope of the Hamming distance relative to the subtype consensus is larger than that relative to the root, while the slope of the RTT regression is even larger. These substantial differences suggest that saturation effects are already strong despite the fact that distances are less than 10%.

The Hamming distance to subtype consensi (green) was computed for HIV-1 subtype B and C sequences, relative to their respective subtype consensus computed from the alignment. The phylogenetic tree was inferred using an IQTree GTR+F+R10 model (Tavare, others, 1986; Yang, 1995; Minh et al., 2020), which was found to be the best model according to the IQTree model finder (Kalyaanamoorthy et al., 2017). For more details, see section M&M 1.2.

Such rapid saturation can arise through rate variation (Soubrier et al., 2012) or heavily skewed site-specific equilibrium frequencies resulting in rapid reversion (Halpern, Bruno, 1998; Hilton, Bloom, 2018; Puller et al., 2020; Ho et al., 2005; Wertheim, Kosakovsky Pond, 2011). We perform a similar analysis on WH data on time scales of 3-10 years to differentiate between the different explanations of rate variation and reversion.

We compute WH evolutionary rates by measuring the divergence over time in Figure 2B. Specifically, we calculate the divergence *d*(*t*) relative to a reference sequence, such as the root of the tree according to:

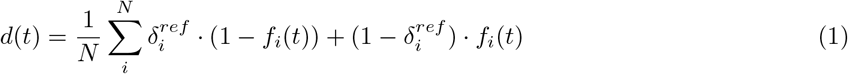

where *N* is the length of the region and *f_i_*(*t*) is the frequency of the founder nucleotide at position *i* and time *t* in the viral population. This founder nucleotide is approximated by the majority nucleotide at the first time point and 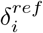 is a Boolean such that 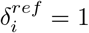 if the founder nucleotide at position *i* is the same as in the reference sequence and 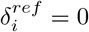 otherwise. More details about the computation of the founder sequence are in section M&M 1.3. When measuring the divergence relative to the founder sequence, we instead use the frequencies of the first sample *t* = *t*_0_ as follows:

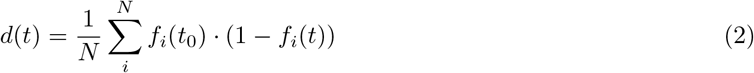

**Figure 2:**
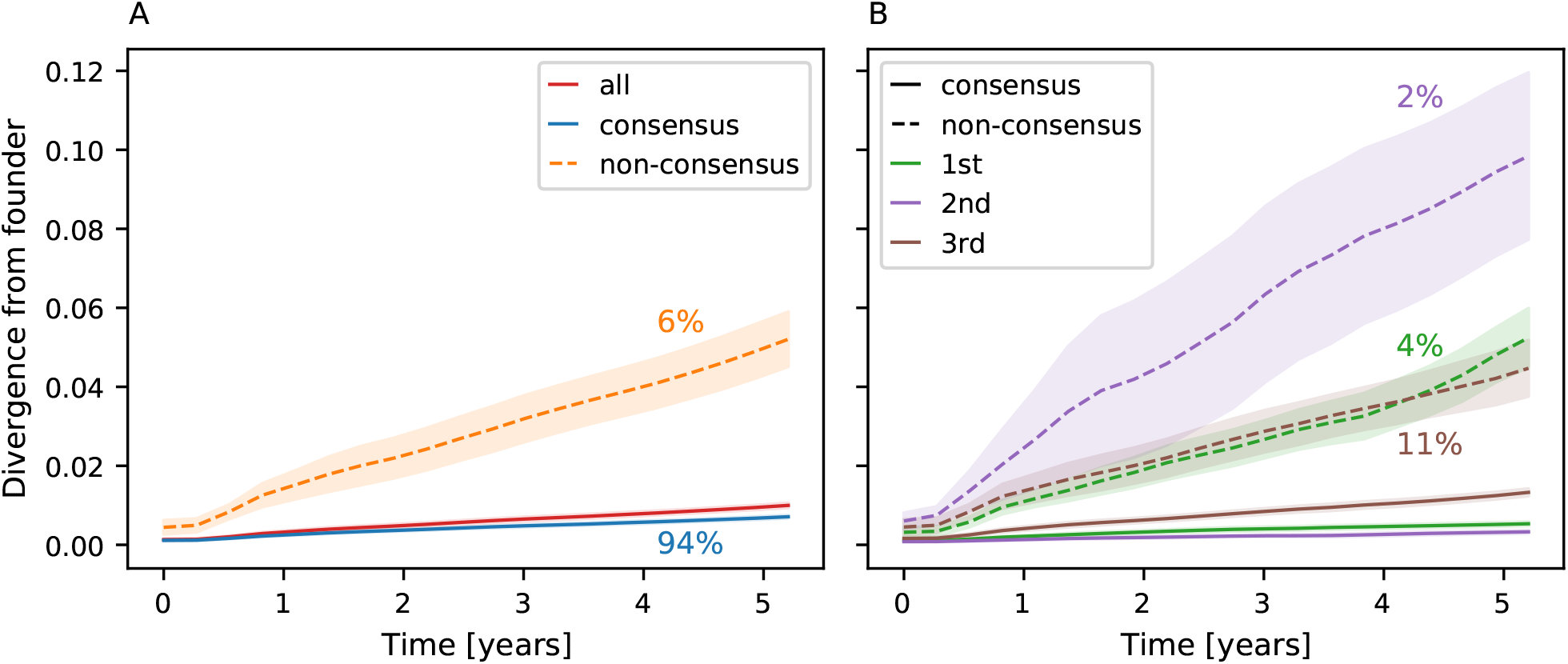
Divergence from founder sequence over time in the *pol* gene. **A**: Divergence to founder overall and split for sites initially in consensus and non-consensus states. The reference used to define consensus and non-consensus sites is the HIV-1 group M consensus. Colored percentages are the fraction of sites corresponding to the related curve. Non-consensus sites represent only 6% of the gene but diverge faster over time. **B**: The same data as that shown in the left panel split between the 1st, 2nd and 3rd codon position. The difference in evolution speed is greatest for nucleotides in the 2nd position, where mutations can not be synonymous. Results for region env and *gag* are shown in Supplementary Figures S3 and S4.

Overall the quantity *d*(*t*) measures the Hamming distance to the reference sequence expected for a randomly picked virus in the viral population of a sample. We then averaged these quantities over all samples and estimated uncertainty by bootstrapping groups of samples from the same patient with replacement. In analogy to the BH analysis, we use the root of the HIV-1 group M tree and subtype consensi as reference sequences, supplemented by the founder sequence of each patient. The results are shown in Figure 1B. The filled areas represent one standard deviation of the bootstrap replicates. For more details about the methodology, see section M&M 1.2. The divergence increases the fastest relative to the founder sequence at approximately (17.1 ± 0.3) · 10^−4^ mutations per site per year, 1.7 times the speed observed from the RTT estimate in the BH data set. Saturation effects play a minor role in the WH evolution speed on the time scale of several years. The rate at which the distance to the root or subtype consensus sequence increases, however, is close to the same estimates in the BH data set. Similar results are found for the env and *gag* regions in Figures S1 and S2.

### Non-consensus sites diverge faster

We next explored the evolution toward and away from consensus within hosts, see Figure 2. Panel 2A shows the WH divergence separately at sites where the founder sequence agrees with the HIV-1 group M consensus and where it differs from it. Filled areas show the standard deviation of the bootstrap estimate. The divergence at sites where the founder sequence differs from the global consensus increases about 7-fold faster than in the rest of the sequence. In the case of the pol gene, about 6% of sites fall into the former category, suggesting that about 1 in 3 changes brings the sequence closer to the HIV-1 root sequence. This is consistent with the 3-fold difference in slope between the divergence relative to the founder or HIV-1 group M root shown in Figure 1B.

This accelerated evolution could be due to (i) reversion to an ancestral state to increase fitness, or (ii) reduced purifying selection at sites with high levels of diversity in global HIV-1 population. In order to differentiate between these possibilities, Figure 2B shows the divergence by codon position. The degree to which reversions are accelerated differs between the 1st, 2nd and 3rd position in a codon. In particular, sites in the 2nd position diverge the fastest when in a non-consensus state, while they diverge the slowest in a consensus state. This is consistent with a particularly strong pressure to revert at the 2nd position, where mutations cannot be synonymous and tend to correspond to more drastic amino acid changes. Accordingly, non-consensus sites in the 2nd position are rare as they represent only 2% of such sites. Similar results are found when using root sequence or subtype consensus sequences as a reference instead. The results for env and *gag* are shown in Figures S3 and S4. These results are consistent with previous observations showing that conserved sites tend to revert more quickly (Zanini et al., 2015) and the notion that selection for reversion is probably driven by the fitness costs of mutations that enabled immune escape in a previous host (Leslie et al., 2004).

Note that more rapid reversion at conserved sites is inconsistent with the hypothesis that rapid reversion is due to an overall elevated evolutionary rate at sites that are often in a non-consensus state. Conserved sites would typically be assigned to low rate categories in phylogenetic inference, while sites that tend to be assigned to high rate categories are those that are diverse and, in this analysis, show the slowest rate of reversion.

### Reversion mutations are positively selected

If a lot of reversions are driven by selection, as the codon position-specific analysis above suggests, effects of selection should be detectable in the dynamics of single nucleotide variants within the host (iSNVs). Specifically, we expect reversion mutations to bear signatures of selective sweeps. We analyzed the frequency trajectories of iSNVs to look for such features. Similar to the previous analysis, we separate all trajectories into reversion and non-reversion groups and compare their evolutionary dynamics in Figure 3. We select trajectories with at least one data point in a frequency interval [*f_min_, f_max_*] for each group. We offset these trajectories in time so that *t* = 0 corresponds to their first data point in the frequency interval and compute the mean frequency of the trajectory group over time. More details about the definition of trajectories and the methodology are in section M&M 1.4 and M&M 1.5. We use the HIV-1 group M consensus sequence as a reference to define reversion mutations, but results are qualitatively similar when using subtype consensus or root sequence as a reference.

**Figure 3:**
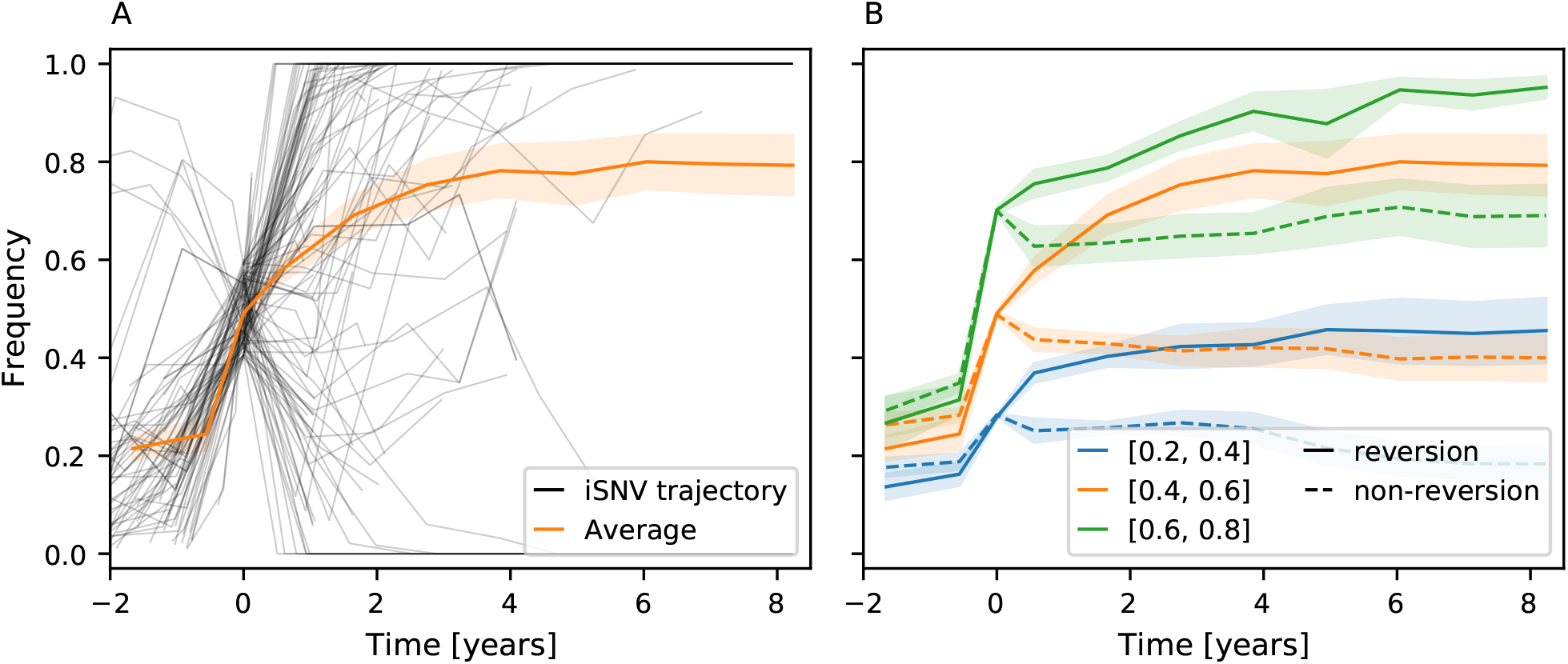
Positive selection on reversion mutations. **A**: Frequency of reversion mutations seen between 0.4 and 0.6 at one time point (offset to be *t* = 0) and their average. **B**: Mean frequency in time for reversion (full lines) and non-reversion mutations (dashed lines) for different frequency windows (colors). Reversion trajectories are strongly selected for as their means increase over time. Non-reversion trajectories evolve close to neutral expectation. The reference sequence used to define reversion mutations is the HIV-1 group M consensus. The solid orange line is the same in both panels.

Figure 3A shows individual trajectories shifted to pass through the frequency interval [0.4, 0.6] at *t* = 0 along with their mean. The mean frequencies for different initial conditions and groups of trajectories are shown in Figure 3B. Since we condition the set of trajectories to start as minor variants and pass through a frequency interval at *t* = 0, we expect that trajectories tend to rise for *t* < 0, as is indeed observed. The dynamics at *t* > 0, i.e. after the time of conditioning, is informative about the selection of the iSNV. We do not expect any consistent trend to rise or fall in frequency for neutral mutations, and the mean frequency should be flat for *t* > 0. Contrary to that, we show in Figure 3B that the frequency of reversion mutations increases on average over time. This suggests that these reversion mutations are beneficial on average and fix preferentially in the population, with probability given by the end point of the curves for each group of trajectories. This finding is consistent with the notion that the HIV-1 consensus sequence approximates a fitness optimum of HIV-1 (Zanini et al., 2017). On the other hand, non-reversion curves are flat or slightly decreasing for *t* > 0, suggesting that such mutations tend to be slightly selected against or are neutral – at least those that reach high frequency in the first place.

We note that the selection for reversion mutations is strongest for the *gag* region and weakest for the env region, see Figure S8 for details. We also observe a difference in selection for synonymous and non-synonymous mutations (Fig. S5), consistent with earlier results (Zanini, Neher, 2013).

### Reversions can explain the rate mismatch

Over longer time scale, the WH reversions we observe will lead to undetected substitutions along branches of the phylogeny whenever a mutation and its corresponding reversion happen on the same branch. When such reversion dynamics are not captured by the substitution models, the evolutionary rate inferred by phylogenetic methods will be too low (Halpern, Bruno, 1998; Hilton, Bloom, 2018; Puller et al., 2020). This bias will contribute to the discrepancy of evolutionary rates within and between hosts.

We quantify the impact of reversion on the BH evolution rate using a WH informed evolutionary model that accounts for the reversion bias highlighted previously. Firstly, we use the TreeTime library (Sagulenko et al., 2018) to define a site-specific general time reversible (GTR) model to reproduce the evolutionary process (Puller et al., 2020). We parametrize the mutation rate from nucleotide *j* to *i* at position *α* as:

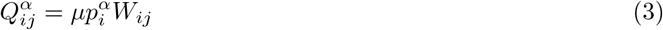

where *μ* is the mean mutation rate per site per year, 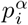 describes the equilibrium probability of finding nucleotide *i* at site *α*, and *W_ij_* accounts for the overall variation in rate between different nucleotide pairs *i* and *j* (e.g. the differences between transitions and transversions). We use *μ* = 17.1 · 10^−4^, the overall WH evolution rate observed in Figure 1B. In this model, the bias for reversion is introduced via the equilibrium frequencies 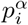. These depend on the genome position *α*, enabling us to skew the frequencies towards the consensus nucleotide at each position. Contrary to common evolutionary models that include rate variation between sites, we keep it constant across positions and vary 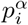 instead. The results are insensitive to Gamma-distributed rate variation with a shape parameter greater than 2. We choose 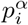 such that the model reproduces the WH rates of reversions and evolution away from consensus. Specifically, we use

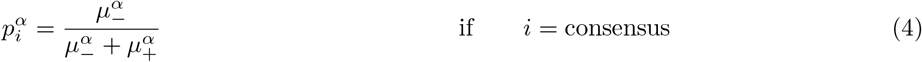

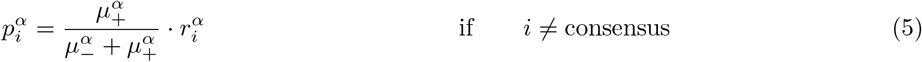

where 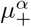 and 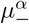 are the consensus and non-consensus evolution rate, respectively, computed from results shown in Figure 2B. These rates are codon position-specific, meaning for every *α* that is a 1st codon position 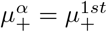 and 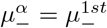. This is analogous for the 2nd and 3rd codon positions. The parameter 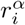 is used to define the relative proportion of non-consensus nucleotides caused by the uneven transition vs. transversion rates. It is chosen so that 85% of the non-consensus nucleotides are the transition from the consensus, while the two transversions contribute 7.5% each. These values were inferred from the BH alignment. Otherwise, this GTR model is purely informed by WH rates.

We then used this model to simulate evolution along a tree and generate a multiple sequence alignment (MSA). This was done using TreeTime by providing the root sequence and tree inferred from the real BH data, as used in Figure 1. We then inferred a tree from the MSA generated using IQTree, as we did for the real data.

Figure 4 compares the diversity of original and generated MSAs and the length of the inferred trees to quantify the impact of reversions on phylogenetic inference. A model that does not account for reversions, i.e. where *p_i_* = 0.25 for *i* ∈ A,C,G,T for all sites, was included for comparison and is referred to as the naive GTR model. Figures 4A and 4B show a comparison of the real and generated MSA characteristics. The MSA generated using our WH-informed GTR model (green) has a similar nucleotide content and distance to root as the real BH data (blue). On the contrary, the naive GTR model that does not take reversions into account (orange) results in a more diverse MSA and overall nucleotide content that is less similar to the BH data.

**Figure 4:**
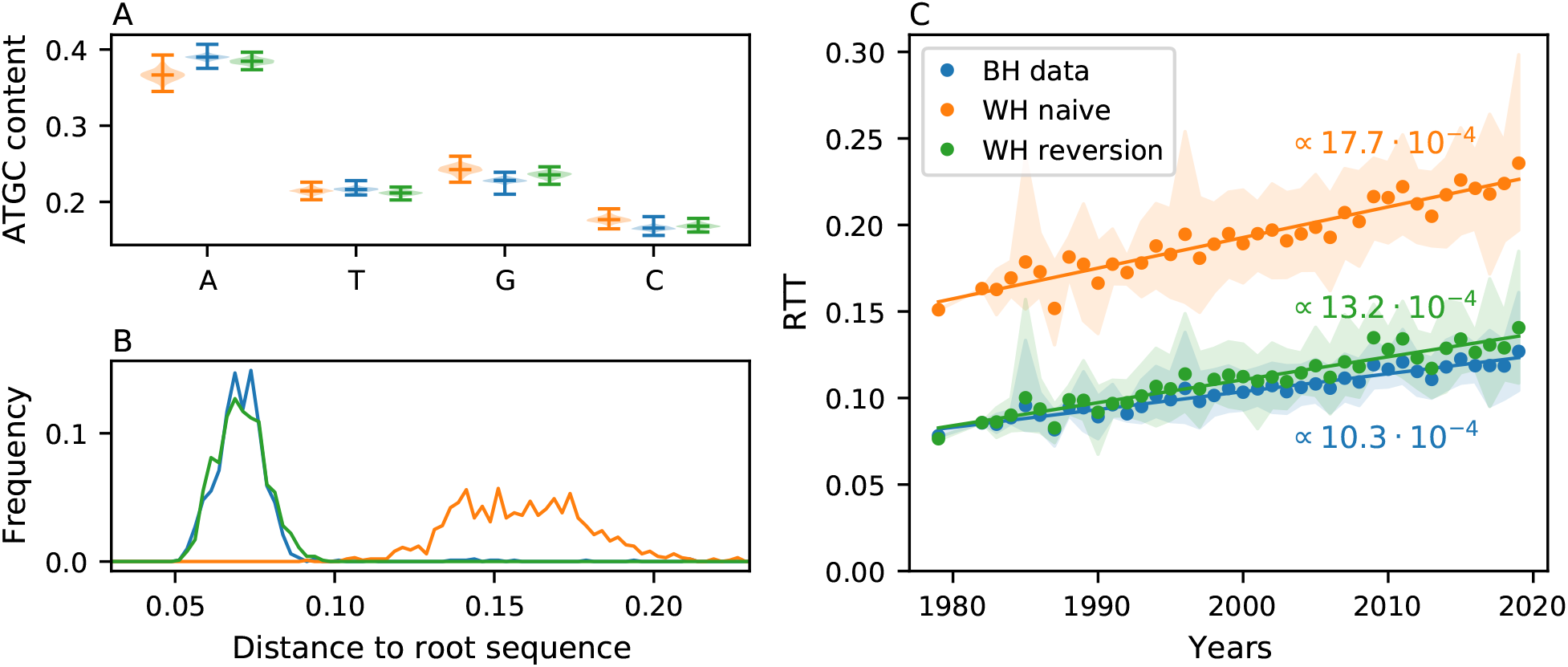
Substitution models that account for reversions can largely explain the rate mismatch. This figure shows the sequence diversity and RTT distances for simulated data generated with a substitution model that account for reversion parametrized by WH data (WH reversion) and a model that does not account for reversion (WH naive) for the pol region. **A**: Violin plot of the nucleotide content for BH and the MSAs generated. The naive model tends to change the nucleotide content, while the model accounting for reversion matches the BH observations better. **B**: Histogram of Hamming distances to the root sequence. The reversion-informed model agrees well with the BH observations, while the naive one drifts too far from the root. **C**: While the RTT distance estimated from data generated by the naive model is correctly estimated (true value 17.1 · 10^−4^), data generated using the model with reversion results in much lower estimates, similar to the rate estimated from BH data. Results for env and *gag* are shown in Supplementary Figures S6 and S7.

A similar behaviour can be observed from the increase of RTT distances over time, as shown in Figure 4C. The evolutionary rate estimated from the RTT regression of the naive GTR model is, as expected, very close to the WH evolution rate of *μ* = 17.1 · 10^−4^ mutation per site per year we input into the model. Our custom GTR model, which uses the same *μ* but accounts for reversions, results in a RTT regression with a slope of 13.2 · 10^−4^. This suggests that a substitution model parametrized by rates and reversion propensity of WH evolution can largely reconcile the discrepancy of rate estimates at different scales.

We find qualitatively similar results for the *gag* and *env* region, see Figures S6 and S7. In the case of *env*, the tree reconstructed from the data generated and subsequent analysis is unreliable, suggesting that reversions attenuate the clock signal significantly.

## Discussion

Evolutionary rate estimates depend strongly on the time scale over which they are measured (Ho et al., 2005; Aiewsakun, Katzourakis, 2016). Here, we explored this effect on the scale of the HIV-1 pandemic, individual subtypes and within hosts and attempt to bridge these scales. We show that differences between WH estimates and rate estimates on the pandemic scale can, to a substantial degree, be explained by a rapid and strong tendency to revert deleterious mutations to their preferred state. These unpreferred states are probably the result of immune selection in a previous host which gradually revert as the host-specific selection pressure is relaxed in future hosts. Microscopically, we thus observe evolutionary dynamics driven by the adaptation to a changing environment. These transiently beneficial escape mutations are generally deleterious on longer time scales, such that the aggregate effect of this dynamic looks like slowly acting purifying selection (Wertheim, Kosakovsky Pond, 2011).

The rate of reversions observed during chronic infection within hosts can explain approximately 60% of the apparent slowdown of evolution for the pol gene of HIV-1. A similar selection for reversion mutations has also been observed during acute infection (Boutwell et al., 2010; Leslie et al., 2004) or the transmission bottleneck (Carlson et al., 2014). Such preferential transmission of consensus-like variants will amplify the overall effect of incomplete reversions during chronic infection. Together, these results suggest that, among the hypotheses proposed to explain the difference in rates (Lythgoe, Fraser, 2012; Redd et al., 2012; Zanini et al., 2015; Leslie et al., 2004), the “adapt and revert” mechanism seem to be the main driver of the slowdown.

Substitution models commonly used to reconstruct phylogenies and infer evolutionary rates do not account for rapid reversions, which require site-specific preferences for different states (Halpern, Bruno, 1998; Hilton, Bloom, 2018; Puller et al., 2020). Such model misspecification is particularly problematic on long branches in the phylogenetic tree, where mutations are masked by their corresponding reversions. Such site-specific saturation effects probably affect deep and shallow parts of the tree to a different degree and deep long branches will suffer more severe distortions.

Rapid reversions are probably essential to conserve global fitness by purging costly immune escape mutations acquired in individuals earlier in the transmission chain (Carlson et al., 2014; Zanini et al., 2017). In addition to reversion, fitness costs of escape mutations can, of course, also be mitigated by compensatory mutations (Crawford et al., 2007; Carlson, Brumme, 2008). Though such compensatory mutations presumably slow down many reversions, we still observe a marked difference in iSNV frequency dynamics towards vs. away from consensus. In addition, compensatory evolution can change the preferred sequence to a new local fitness maximum to which mutations revert, adding an additional slow time scale to the evolutionary process. We expect the preferred sequence to slowly drift on time scales much longer than the typical serial interval. This effect has been observed in deep mutational scanning experiments in influenza viruses (Hilton, Bloom, 2018; Doud et al., 2015).

The star-like diversification of HIV-1 into multiple subtypes gives a clear notion of a consensus sequence that can be used to approximate a putative fitness peak towards which reversions occur. In other viruses, for example, influenza A viruses, the ladder-like or otherwise structured phylogenies do not allow a straightforward definition of a consensus sequence. Nevertheless, it is possible that adaptation to a changing immunity landscape and reversions contribute with a similar magnitude to sequence turnover.

## Acknowledgements

We gratefully acknowledge stimulating discussions with Pierre Barrat-Charlaix and Marco Molari. This work was funded by the core funding of the University of Basel, the SNF (310030_188547) and the PhD Fellowship Program of the Biozentrum (to VD).

## 1 Material and Methods

### 1.1 Data set and filtering steps

#### Between host data sets

Our BH data sets come from the Los Alamos National Laboratory HIV databases. All HIV-1 group M sequences containing 0% problematic nucleotides in the region and an exact sampling date were downloaded for each region. Subtype O and subtype N sequences were filtered out. Only one sequence was kept per patient. Data were downloaded on July 14, 2021. This gave us a total of 6649 sequences for *pol*, 15034 for *env* and 8948 for *gag*.

Regarding each genomic region, we subsampled the data set to have 1000 sequences in each case, with the same number of sequences for each year where sequences were available (except for early years where fewer sequences were available). We then performed an MSA, including the HXB2 sequence, using MAFFT (Katoh, Standley, 2013) and the Nextstrain framework (Huddleston et al., 2021). Insertions relative to the reference HXB2 sequence were removed. We removed all positions of the alignment where more than 10% sequences have a gap as the alignment can be unreliable in such positions. The alignment for the *pol*, *env* and *gag* regions are the data sets used for our BH analyses. See section Code and data availability for access to the data sets.

#### Within-host data sets

Our WH analysis leverages the time resolution of the HIVEVO data set (Zanini et al., 2015). This data set is freely available with tools made available to facilitate the analysis. We use these tools to obtain a three-dimensional matrix of nucleotide frequencies for each patient. The three axes of these tables are the HIV-1 genome position, the nucleotide and the time since infection of the sample. Each entry in these matrices gives the frequency of a given nucleotide at a given position on the genome at this time point, relative to the total intra-patient HIV-1 population. These matrices form our WH data set. We excluded patient p7 and *p10* from our analysis as their samples were very uneven in time.

Estimates of nucleotide frequencies are unbiased in the [0.1,0.9] range, while coverage and depth are globally sufficient (Zanini et al., 2016). We applied several filtering steps prior to analysis to avoid biases in our results. We, firstly, masked data points with sequencing coverage inferior to 100 and/or where the depth was low. We also removed genome positions that were not mapped to the consensus sequence and/or seen to be too often gapped in the MSA of BH sequences. The alignment and mapping of such sites can be unreliable, thus, we removed them from our analysis. This filtering procedure is mainly relevant for the *env* gene, which is the region with the most noise.

### 1.2 Distance and divergence over time

The first result section gives an overview of the method used to compute the distance and divergence over time in Figures 1, S1 and S2. Additional details are given below.

Hamming distances were computed by counting the number of sites that do not match the reference sequence for each sequence in the data set. We then divide this number by the length of the sequence to obtain the relative distance to the reference. Hamming distances were computed using three reference sequences. The first is the root sequence of the tree. The tree was inferred using the IQTree GTR+F+R10 model (Minh et al., 2020), while the root sequence was computed using TreeTime ancestral reconstruction on this tree (Sagulenko et al., 2018). We chose to use the root sequence instead of the consensus sequence of the alignment in Figures 1 and 4 to avoid biases due to overrepresentation of subgroup B and C sequences. The second and third reference sequence are subgroup B and subgroup C consensus sequences. See section M&M 1.3 for details about the computation of consensus and founder sequences. We performed an average of the distance computed for subtype B and C sequences relative to their consensus to compute the Hamming distances to subtype consensus. The average was weighted by the relative number of each subtype sequence in each year.

The RTT distances shown in Figures 1, 4, S1 S2 S6 and S7 are computed directly from the tree generated via IQTree. Such distances were computed for every leaf of the tree (i.e. every sequence in our data set) and then averaged for sequences sampled in the same year for visualization purposes. Taking into account the phylogenetic information allows one to detect some mutations that would occur back and forth along the tree. Consequently, the estimates of the RTT distance are higher than the Hamming distance ones.

### 1.3 Consensus and founder sequence

Consensus sequences were computed from our BH data sets. We computed three consensus sequences for each region studied. First is the HIV-1 group M global consensus, which is the majority nucleotide of the alignment at each position. Second and third are the subtype B and subtype C consensus sequences. These were computed in the same way, using a subset of the alignment that contains only the sequences of the subtype in question.

The founder sequence is an approximation of the sequence of the virus at the time of infection in a patient. They are computed from our WH data set for each patient separately. The founder sequence is the majority nucleotide in each position from the first sample of each patient. In this sense, it is the consensus sequence obtained from the first sample of each patient. For most patients in our data set, the first sample is taken at approximately 90 days after infection and no data is available on the early phase of infection. Consequently, the founder sequence computed is an approximation of the original virus.

### 1.4 Trajectory extraction and metadata

A trajectory is a sequence of nucleotide frequencies and associated time. Each trajectory corresponds to one genome position and one nucleotide only. We extracted trajectories from our WH data set according to several criterion. Firstly, every trajectory is extinct before the first point, i.e. we consider only new mutations. This is to avoid biases that could be due to immune interaction existing already. Secondly, frequencies are between 0.01 and 0.99 at all time points. The trajectory is considered extinct if it is below 0.01, and fixed if above 0.99. Lastly, we apply a mask to data points according to what is shown in section M&M 1.1. Trajectories that have their first and/or last point masked are removed from the analysis.

Every trajectory extracted according to the criterion above is coupled with its metadata. This contains all the relevant information, such as whether the mutation is a reversion or not and whether it fixed or was lost. This information is used to create subgroups of trajectories. From these subgroups, one can study the impact of a trait associated with a mutation for WH evolution, as shown in Figure 3 for reversion and non-reversion trajectories.

### 1.5 Mean frequency in time

While looking at divergence values informs us on the global evolution of the WH population, it cannot tell us whether the mutations we see on non-consensus sites are actually reversions to the consensus state or simply mutations to another nucleotide. This motivated us to look directly at the evolution of new mutations independently by observing the frequency trajectories in time. Trajectories were extracted and filtered according to sections M&M 1.1 and M&M 1.4. Despite these filtering steps, our data is inherently biased towards small and/or low-frequency trajectories which are more common. In order to alleviate this bias, we compare reversion and non-reversion trajectories in the same manner. Accordingly, the resulting signal can be attributed to the effect of being a reversion (or not).

Due to the limited number of trajectories available and the often lack of information about trajectory fixation, for example, because it is still active at the last sample, the probability of fixation plots were not adequate for our analysis. We, thus, decided to pay attention to the evolution of the mean frequency in time of groups of trajectories. Trajectories were grouped in frequency bins, as described in the main text, to avoid bias towards positively or negatively selected trajectories.

We then created time bins of 400 days from 600 days before up to 3000 days after a trajectory is seen in a frequency window. We compute the average frequency of all trajectories belonging to the same group in each time bin. A trajectory contributes its current frequency if a data point is available at this time, and does not contribute if no data is available in that time bin. Trajectories that fixed in the population contribute with a frequency of *f* = 1 to time bins subsequent to their fixation. Similarly, lost trajectories contribute *f* = 0 to time bins subsequent to their disappearance in the viral population. Trajectories that are still active after their last data point (because the study stopped before it could fix or be lost) contribute the frequency of their last data point to the following time bins.

## Code and data availability

The code and data used for the analysis can be found at https://github.com/neherlab/HIVEV0_reversion. Due to issues with the data sets’ size, only intermediate BH and WH data files in a compressed format are found in the github folder. A link to the full data set is available there. Scripts are present to reproduce the results shown in this paper.

## Supplementary Materials

**Supp. Fig. S1:**
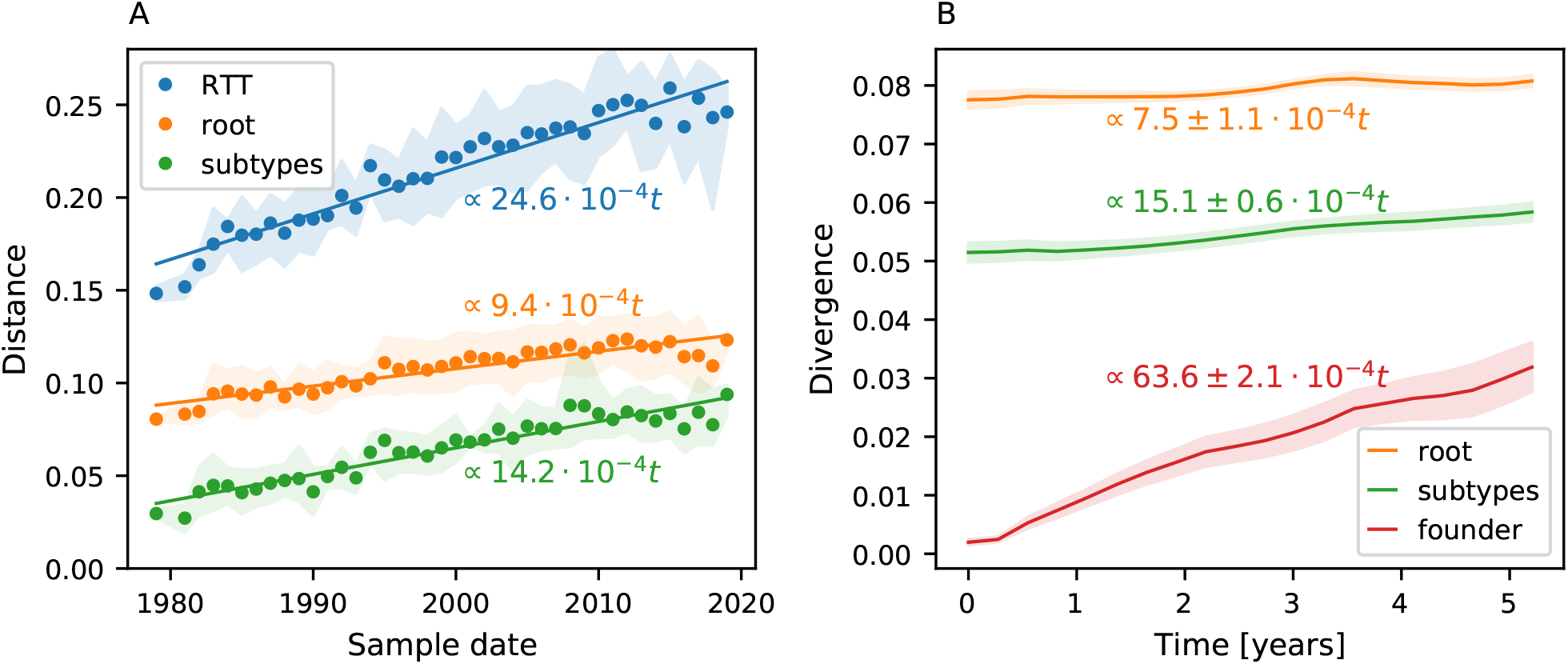
Same as Figure 1 for the *env* gene. The Y axes are not shared in this case. The *env* gene behaves similarly to *pol* and *gag* but the difference in the rate observed is higher. This is consistent with the fact that *env* mutates faster overall, which would also lead to more reversions.

**Supp. Fig. S2:**
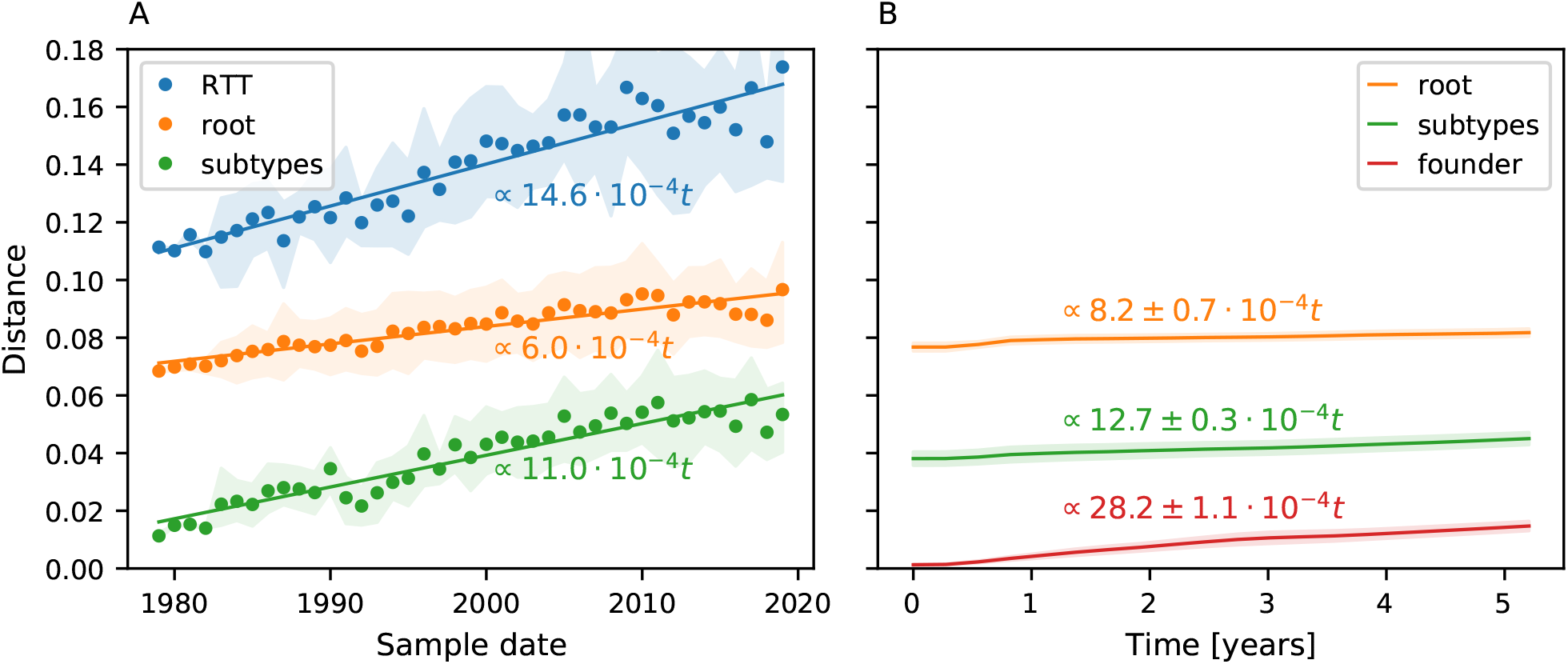
Same as Figure 1 for the *gag* gene.

**Supp. Fig. S3:**
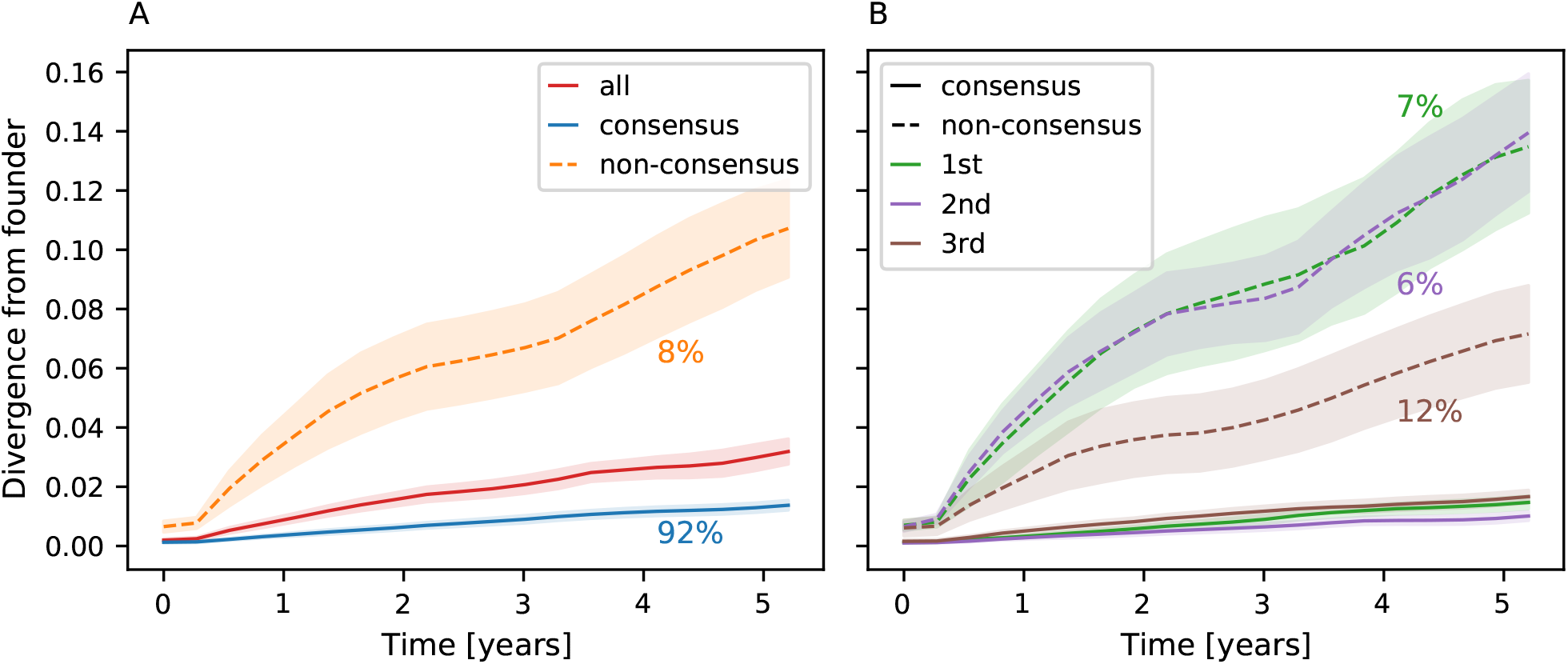
Same as Figure 2 for the *env* gene.

**Supp. Fig. S4:**
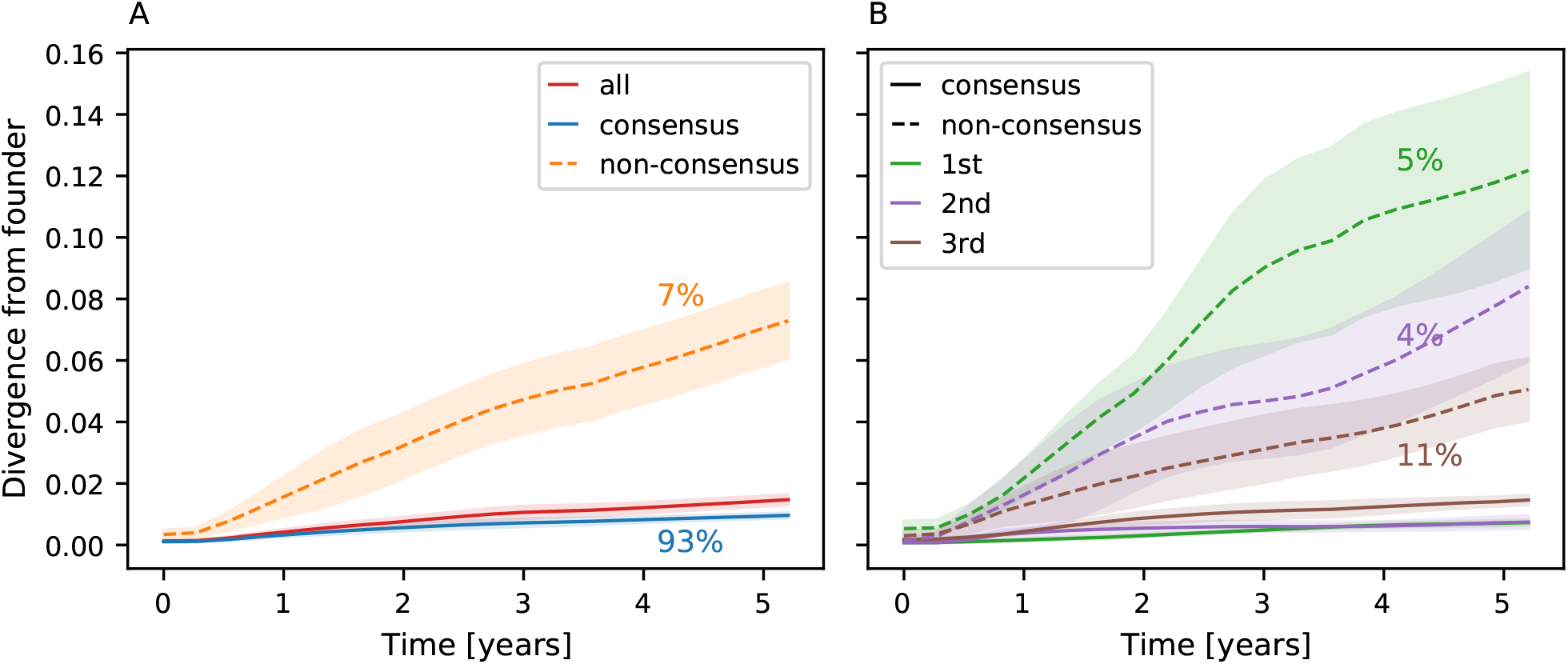
Same as Figure 2 for the *gag* gene.

**Supp. Fig. S5:**
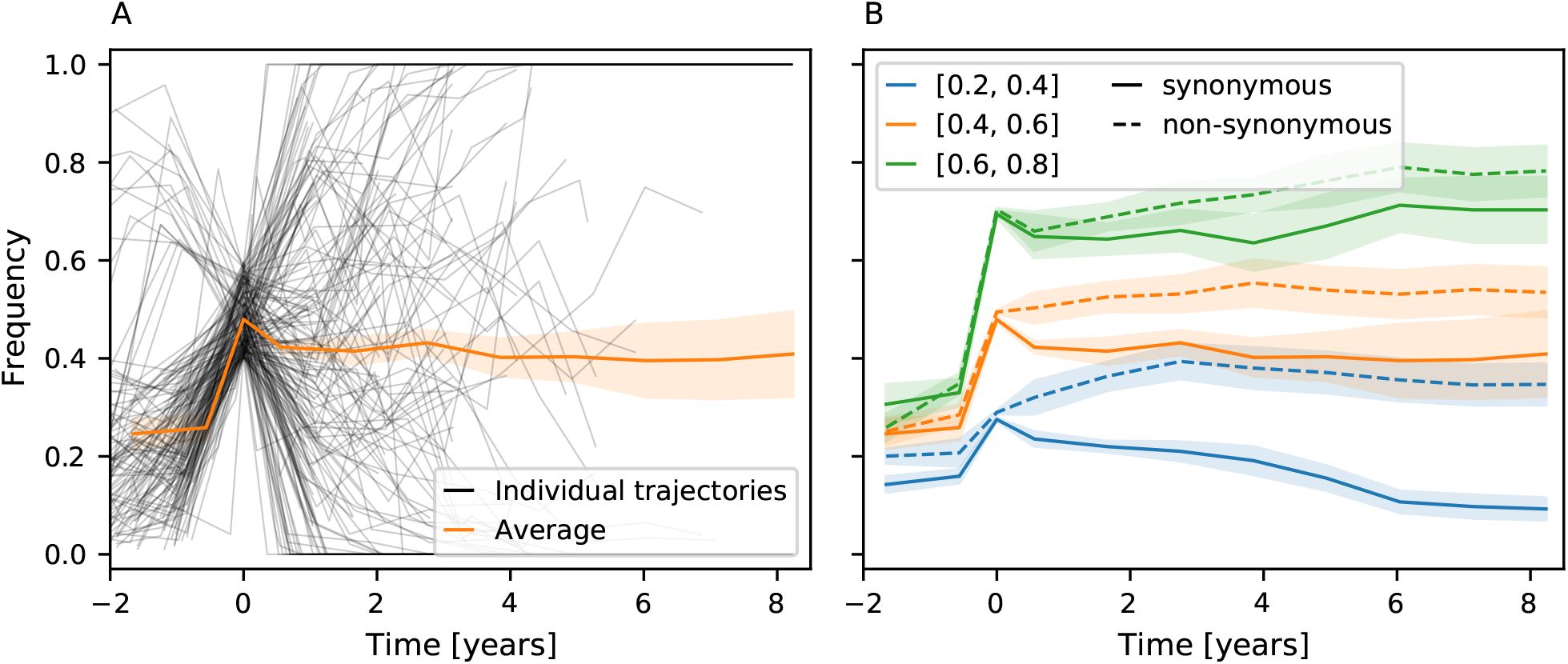
Same as Figure 3 but for synonymous and non-synonymous trajectories. Overall synonymous mutations are selected against and non-synonymous mutations seem to be selected for, but the effect is smaller than what we see for reversions and non-reversions in Figure 3.

**Supp. Fig. S 6:**
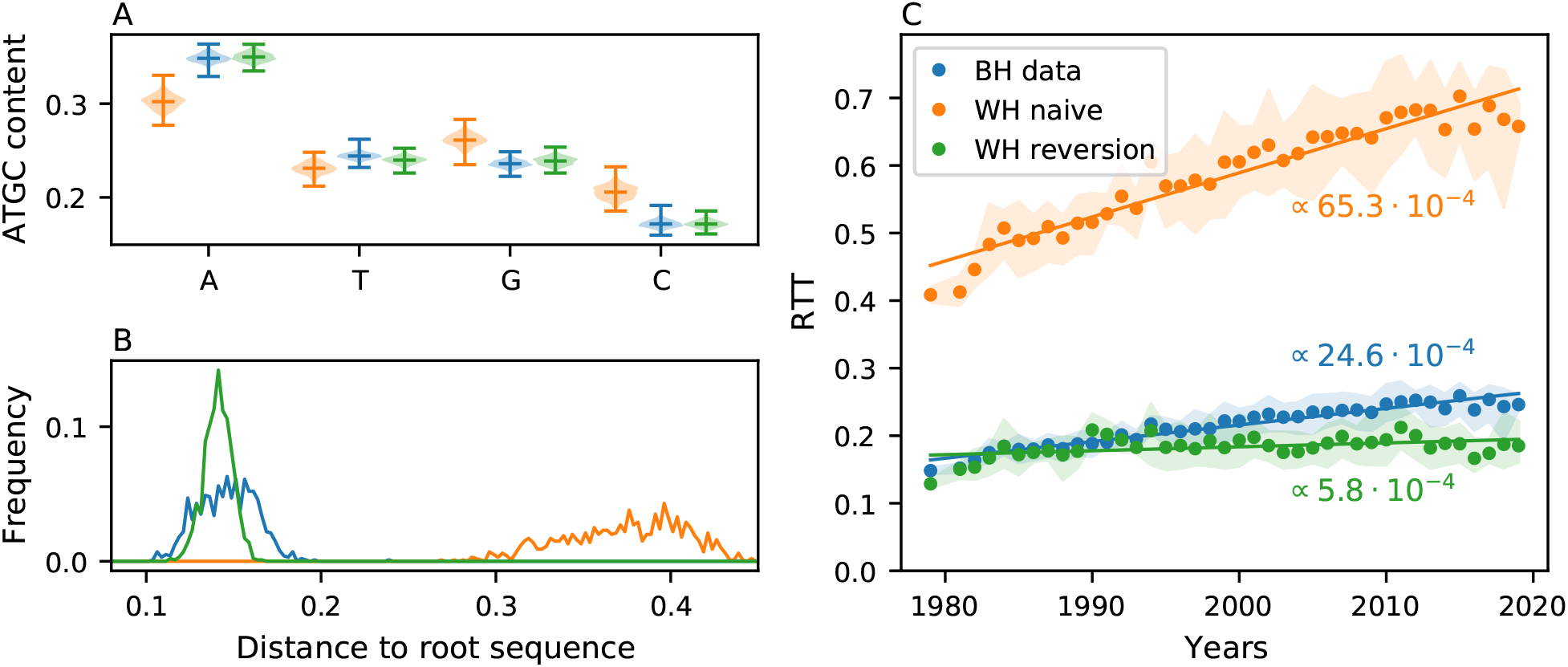
Same as Figure 4 for the *env* gene. The WH mutation rate in this region is so high that the reversion model attenuates most of the clock signal, which leads the tree reconstruction to fail.

**Supp. Fig. S7:**
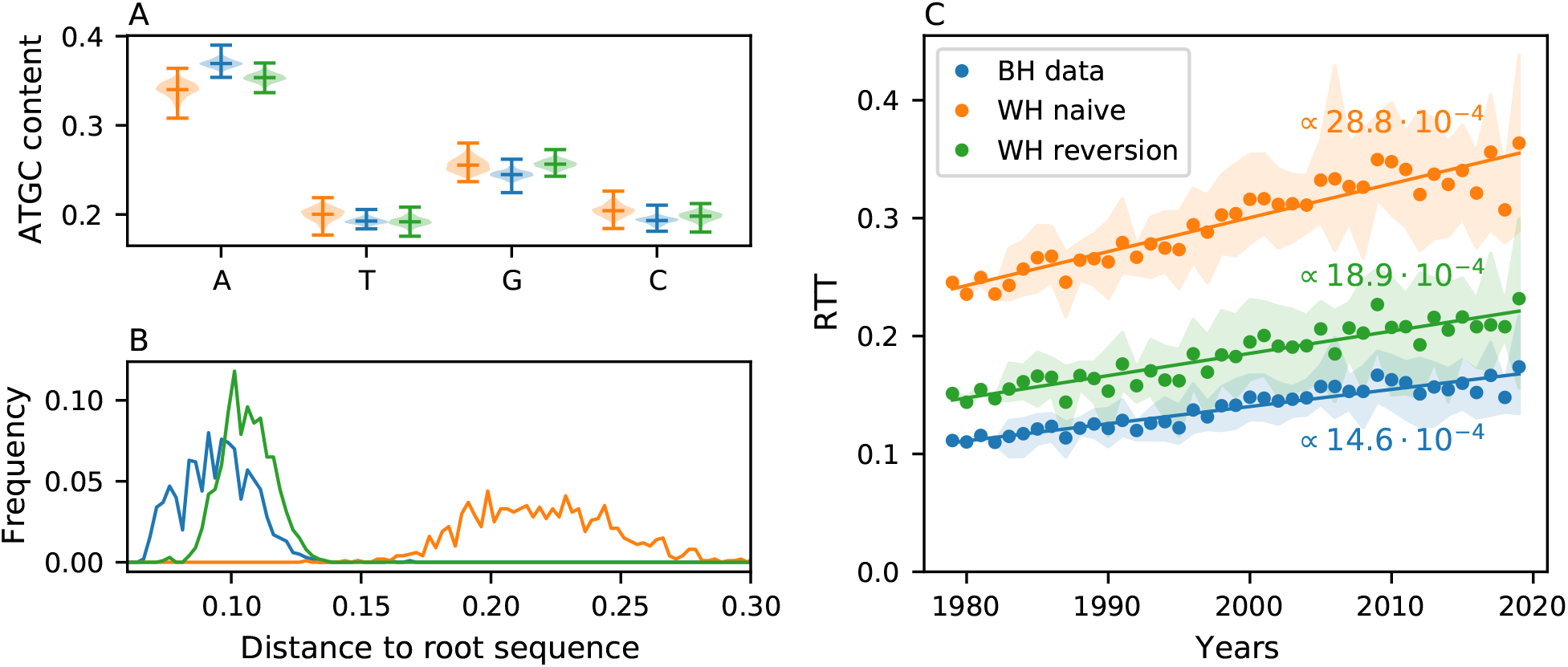
Same as Figure 4 for the *gag* gene.

**Supp. Fig. S 8:**
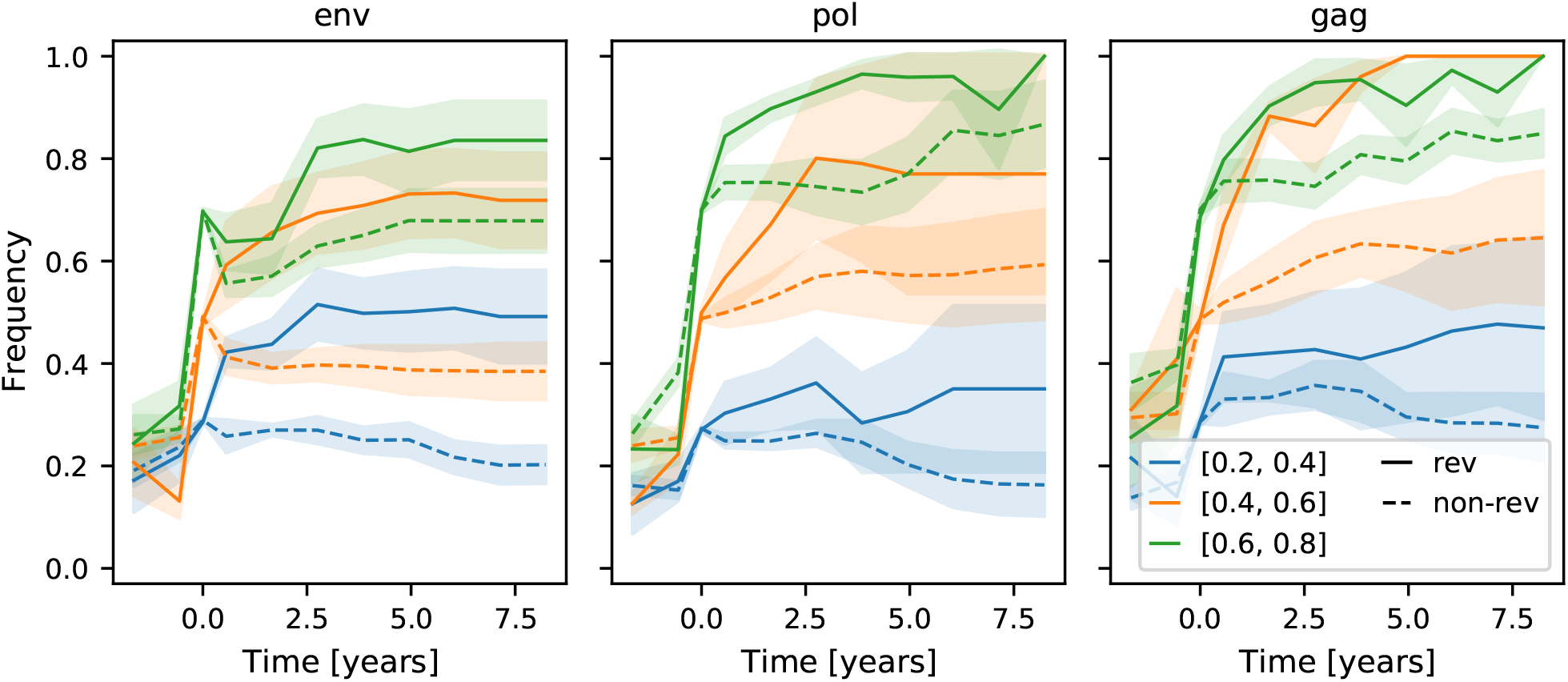
Same as Figure 3B split per region. Selection for reversion is strongest in the *gag* region and weakest in the *env* region.

## Notes

### Competing Interest Statement

The authors have declared no competing interest.

